# Predicting non-coding RNA function using Artificial Intelligence

**DOI:** 10.1101/2024.12.30.630736

**Authors:** David da Costa Correia, Francisco M. Couto, Hugo Martiniano

**Affiliations:** Departamento de Promoção da Saúde e Prevenção de Doenças não Transmissiveis, Instituto Nacional de Saúde Doutor Ricardo Jorge, Avenida Padre Cruz, Lisboa, 1649-016, Portugal; BioISI - Biosystems and Integrative Sciences Institute, Faculdade de Ciências da Universidade de Lisboa, Campo Grande, Lisboa, 1749-016, Portugal; Departamento de Informática, Faculdade de Ciências da Universidade de Lisboa, Campo Grande, Lisboa, 1749-016, Portugal; LASIGE, Faculdade de Ciências da Universidade de Lisboa, Campo Grande, Lisboa, 1749-016, Portugal

**Keywords:** non-coding RNAs, Relation Extraction, Text Mining, Distant Supervision, Large Language Models

## Abstract

Non-coding RNAs (ncRNAs) represent the majority of human gene products, and are involved in various important biological processes, being considered relevant disease biomarkers and therapeutic agents. However, information about these biomolecules remains sparsely distributed, mostly in the form of scientific research articles. It is then of pivotal importance to aggregate and summarize the existing information.

Natural Language Processing (NLP) methods applied to text mining can be used to generate collections of annotated sentences expressing relations between entities, called relational corpora.

In this work we developed a text mining pipeline to generate a ncRNA-phenotype relational corpus (ncoRP) using Distant Supervision Relation Extraction (DSRE), comprising 21,608 annotated articles, 2,835 unique ncRNAs, 1,118 unique phenotypes and 35,295 unique relations, with a precision of 0.761 and F1-score of 0.593, calculated through human validation. DSRE methods require a set of pre-documented relations to function, as such, a high-fidelity ncRNA-phenotype relation dataset, consisting of 214,300 unique relations, was created by the aggregation of five comprehensive ncRNA-disease functional annotation databases. Then, both ncoRP and the relation dataset represent important contributions towards solving the problem with the sparseness of information about ncRNAs.

Large Language Models (LLMs) are an emergent type of language model, showing great capabilities in general task-solving through text generation, without the requirement of fine-tuning with large datasets. In this work, a LLM RE methodology is proposed and evaluated, achieving an F1-score of 0.978 by combining the RE task with a preceding sentence filtering task and applying prompting principles such as in-context learning and Chain-of-Thought self-explanation.

## 1. Introduction

Non-coding RNA (ncRNA) is a broad term that encompasses all RNA molecules that do not encode proteins. ncRNAs are involved in a wide range of cellular processes, such as protein synthesis, regulation of gene product expression and other biological processes [1]. Non-coding regions constitute the majority of the human genome - according to the HUGO Gene Naming Committee (HGNC^1^), there are 19,393 documented protein-coding genes and 25,641 documented non-coding genes, whereas 9,307 of them represent ncRNAs.

The most trivial ncRNAs that come to mind might be ribosomal RNA (rRNA) and amino acid transport RNA (tRNA), both directly involved in the translation step of protein synthesis [1]. Other types of ncRNA, depending on their size, can be classified as micro RNA (miRNA), and small/long non-coding RNA (sncRNA or lncRNA) [1, 2].

MiRNAs and lncRNAs play important roles of cytoplasmic regulation [3]. The former is involved in signaling for repression or degradation of messenger RNA (mRNA), while the latter interacts with various regulatory elements of transcription, such as enhancers and even miRNAs [1]. Moreover, lncRNAs also take part in other regulatory processes, such as X chromosome silencing, genomic imprinting and chromatin modification [2]. Other less prominent types of ncRNA in research literature are small nucleolar RNA (snoRNA), circular RNA (circRNA), small interfering RNA (siRNA) and PIWI-interacting RNA (piRNA) [4]. With this wide range of functions, it is not surprising that dysregulation of ncRNA expression could be a cause of disease and its aggravation [5]. Consequently, ncRNAs can serve not only as diagnosis biomarkers, but also as therapeutic agents [1, 2]. This latter application shows promise in the sense that, due to their specificity to certain mRNAs or proteins, ncRNAs can alter these molecules’ function and structure. Thus, RNA therapies could increase the number of druggable targets, enabling and enhancing other therapies [4].

However, disproportionately to their apparent importance, ncRNAs still remain poorly studied and documented, mainly due to their sheer complexity. It is difficult to understand unique function in the midst of the complex networks and biological pathways that ncRNAs are inserted [4]. Most of the times specific biological studies are required, which are expensive and time consuming [2]. In addition and in consequence of this, even though there are ncRNA databases there are only a few dedicated to functional annotation [2], with the majority of information on ncRNAs still being dispersed in the form of scientific research articles, thus making it essentially impossible to be aware of its full extent by manual interpretation [6, 7]. Considering all this, it is of pivotal importance to aggregate and summarize the available information in a way that enables easy comprehension of existing associations, thus avoiding wasteful repetition of studies which recursively aggravates the problem.

Natural Language Processing (NLP) is the sub-field of Artificial Intelligence (AI) that enables computers to automatically understand human language in its many nuances, and encompasses Text Mining (TM) tasks such as Information Extraction (IE) and summarization [8, 9]. Oftentimes, the first step in TM pipelines is the Entity Recognition and Linking (ERL), responsible for identifying named-entities in text - Named-Entity Recognition (NER) - and subsequently linking them to a knowledge-base entry - Named-Entity Linking (NEL). Then, Relation Extraction (RE) is applied to identify relations between the named entities. A simple starting point algorithm for this task is Distant Supervision Relation Extraction (DSRE), based on the simple application of the following principle: If a relation between A and B is pre-documented in a knowledge base, then a sentence mentioning both A and B expresses that rela-*tion*.

One application of these techniques is in the creation of relational corpora, which are collections of sentences annotated with found relations between entities there mentioned [10]. Thus, the creation of a corpus containing ncRNA-phenotype associations could be a step towards solving the existing problems with the sparseness of information on this topic. Moreover, relational corpora can be further used to develop and train Machine Learning (ML) models to perform various NLP tasks.

Large Language Model (LLM) research is currently on the rise due to these models’ advanced understanding and generation of language [11]. These models also show good performance on general task solving without requiring task-specific fine tuning, unlike traditional state-of-the-art NLP model architectures. As so, further exploration on the potential of LLMs for this kind of problem may be of worth.

## 2. Tools and Methods

This work can be divided into three phases: 1) the creation of a ncRNA-phenotype relation dataset, which directly supports the 2) the creation of a ncRNA-phenotype corpus - ncoRP - with DSRE and 3) the development of a LLM-based Relation Extraction (RE) methodology, using the validated subset of ncoRP to evaluate performance. All the described pipelines were implemented using the Python programming language.

### 2.1. ncRNA-Phenotype Relation Dataset

In the first phase, five ncRNA-disease functional annotation databases (Human microRNA Disease Database [12], lncRNA-Disease Database [13], ncrPheno [14], RIscoper [15] and RNA-Disease [16]) are processed and aggregated in a ncRNA-phenotype relation dataset.

#### 2.1.1. ERL of phenotypes in disease names/descriptions

The selected databases (with the exception of RNADisease) do not provide an uniform representation of phenotypes, often simply using disease names, abbreviations or acronyms. As so, ERL of phenotypes solves this problem by i) assigning IDs to redundant descriptions (i.e. “gastric cancer” and “stomach neoplasm”) and ii) rejecting uninformative or ambiguous descriptions.

To do this, a pipeline using SentenceTransformers^2^ and Facebook AI Similarity Search (FAISS)^3^ was implemented. The first, SentenceTransformers, is a Python module that enables the training and usage of text (and image) embedding BERT-based models, offering a collection of already pre-trained and ready-to-use models [17]. The second, FAISS, is a library (with a Python wrapper) that enables efficient similarity search of vectors (as are embeddings), using different distance metrics such as Euclidean, dot product and Cosine [18].

Therefore by using SentenceTransformers to create a collection of embeddings from the Human Phenotype Ontology (HPO) [19] in which each term name (and respective synonyms) is converted to an embedding associated with its HPO ID, it is possible to find the HPO term embedding most similar to an embedded disease description using FAISS. A distance threshold can also be set in order to filter descriptions that may be too dissimilar from any HPO term. The SentenceTransformers model used was “all-MiniLM-L6-v2”, and a maximum euclidean distance threshold of 0.5 was imposed.

This method was manually evaluated in a random sample of 900 unique disease descriptions (from which 450 linked to HPO IDs), representing about 10% of the 8,772 total unique disease descriptions, yielding a precision of 0.973, a recall of 0.883 and a F1-score of 0.926 and resulting in 4,954 unique disease descriptions linked with HPO terms.

#### 2.1.2. NEL of ncRNAs

In order to ensure that all ncRNAs names registered in the dataset represent actual ncRNA molecules, these were linked to RNACentral [20] IDs. The IDs were assigned through rule-based matching of the ncRNA name to the RNACentral mappings file. NcRNA names that failed to link to an ID were excluded from the dataset, remaining 5,215 unique ncRNA names out of the initial 73,770.

#### 2.1.3. Propagation of relations to phenotype ancestors and ncRNA aliases

To maximize the size of the dataset, and so, the number of positive labelled sentences: 1) relations involving a ncRNA were propagated to all its documented alias names and 2) relations involving a phenotype were propagated to all the HPO ancestors of said phenotype. To obtain the aliases for each ncRNA, the custom downloads tool of HGNC4 was used to download a file containing the “Approved symbol”, “Previous symbols” and “Alias symbols”, filtering to obtain only entries representing the “non-coding RNA” locus group. Then, for each relation, if the ncRNA had aliases (i.e. “Previous symbols” and/or “Alias symbols”), the relation was propagated to each of them.

In the case of phenotype ancestors, for example, relations involving “Stomach cancer”, are propagated to “Neoplasm of the stomach” and to all its other HPO ancestors until “Phenotypic abnormality”, exclusively. This cutoff is applied because i) the term “Phenotypic abnormality” is too general to have medical significance and ii) every original relation would be propagated to it, making it redundant.

### 2.2. ncRNA-Phenotype Corpus - ncoRP

In the second phase, a text mining pipeline is implemented to 1) obtain scientific research articles that have information on ncRNAs, 2) extract sentences in these articles that mention both ncRNAs and clinically significant phenotypes and 3) identify relations between these mentioned entities. In the end, a human validation of a subset of the corpus was performed.

#### 2.2.1. Article Download and Processing

In the first step, the evidence articles from the ncRNA-phenotype relation dataset are downloaded using NCBI’s E-utilities and then processed to be split into paragraphs and then sentences, with paragraphs shorter than 500 characters and sentences shorter than 50 characters being rejected in order to avoid scientifically uninformative text (i.e. author names, acknowledgment section text, etc).

#### 2.2.2. ERL of ncRNAs and phenotypes in sentences

Then, in the second step, MER [21] is used to identify mentions to ncRNAs and phenotypes in these sentences. MER is a minimal dictionary lookup named-entity recognition and linking tool, that requires only a small set of text files (called a lexicon), containing the names and IDs of entities. Merpy is its Python interface, enabling easy integration with the rest of the pipeline and offering lexicon managing functions. A small modification had to be made for Merpy to work with entity names containing dashes (“-”), as is the case of ncRNAs. The first step was to create the ncRNA and phenotype lexicons. For the former, the ncRNA names and IDs were obtained from the RNACentral mappings file, on which a filter was applied to remove generic names such as “non-coding RNA” and names that would cause the wrong recognition of small words or letters, such as “TR” and “E”. For the latter, the HPO terms and the synonym names of each term were added to the lexicon. Only the HPO terms descendant of “Phenotypic abnormality” were considered, as done in [22], in order to exclude terms with no clinical significance such as “localized” and “frequency”. Then for each sentence,

Merpy was executed twice, to recognize both the ncRNAs and phenotypes there mentioned, with sentences not mentioning at least one ncRNA-phenotype pair being rejected.

#### 2.2.3. Distant Supervision Relation Extraction of ncRNA-phenotype relations

The final stage in the creation of the corpus was the DSRE. To do this, for each sentence, if a row reflecting the relation between the ncRNA-phenotype pair, for the article where the sentence originates was found in the relation dataset, then the sentence was labelled as a *Positive*, otherwise, it was labelled as a *Negative*.

#### 2.2.4. Validation

To be aware of the general quality of ncoRP, a human validation was performed. To do that, random samples of 40 sentences (20 positives and 20 negatives) each were distributed among expert curators to be evaluated. In order to evaluate the fidelity of the curators’ validation, out of the 40 sentences sent to each curator, 20 overlap sentences were the same for every curator, which enabled to compute the Fleiss’ Kappa to evaluate curator agreement. For each sentence, the curator had to choose one of three possible evaluations: Correct (when the sentence label correctly reflected the relation between the entities); Incorrect (on the contrary); or Uncertain (when due to ambiguity, error or any other reason, it was hard to attribute one of the other two evaluations).

### 2.3. LLM Relation Extraction

In the end, the potential of LLMs for RE tasks is evaluated. To do that, an Ollama-based framework is implemented to easen the handling and evaluation of LLMs in binary classification tasks (as is RE). Ollama is an open-source framework, available (not exclusively) as a Python library, that enables easy download and interaction with various pre-trained LLMs. The following LLMs, all available through Ollama, were tested: Llama3 (8B and 70B), Phi3 (3B and 14B), Mixtral (8×7B and 8×22B), Gemma (7B) and Gemma2 (9B).

With the validation of ncoRP, by the direct analysis of the curators’ evaluations, it was possible to produce a ground-truth dataset containing high confidence labels. Then, to evaluate the performance of a model when predicting an instance, its prediction is compared to the high confidence label for that instance. This groundtruth dataset was divided into three subsets: 1) Train, used to produce examples for in-context learning; 2) Test, used to select the best performing method (combining a model, prompt and number of examples) and Validation, used as a final evaluation of the method deemed the best. This was done in order to i) ensure that the iterative method design/test process was not contaminated by prediction bias from in-context examples and to ii) confirm that the performance of the found best method was not due to test instance bias.

The process used to design a prompt that instructs a LLM for RE while ensuring the outputs can be automatically processed can be generally expressed as the iteration of the following steps:

1. Design a prompt that conveys the RE task and output format
2. Design a parsing function to handle the response
3. Observe and evaluate LLM responses
4. Repeat, making adjustments in 1. and 2.

In this stage, it was noticed that the inclusion of incontext (input-output) examples in the prompt drastically reduced the number of invalid model responses (see Figure 2), making them essential. Furthermore, the self-explanation Chain-of-Thought principle (prompting the model to explain the reasoning behind its answer) was also included, as it has shown to increase performance, but also give insight on the reasoning behind a prediction [23], which assisted in the prompt design. Once the final prompt (Figure 1) and parsing function combination was achieved, the impact of the number of shots on performance was further analyzed in two separate methodologies: 1) single prompt for Relation Extraction only and 2) a prompt for Relation Extraction preceded by a prompt for Uncertain Filtering.

**Figure 1:**
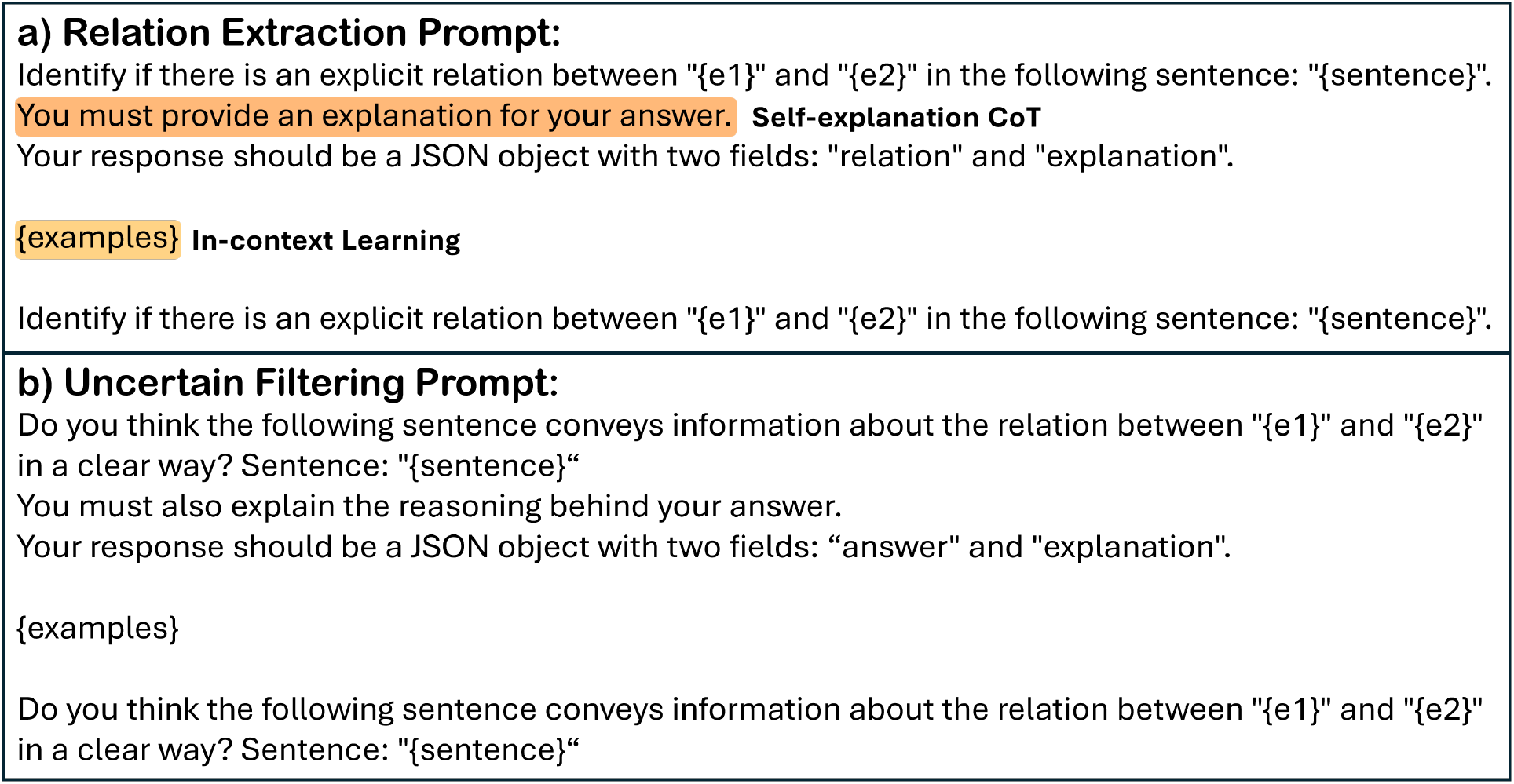
Prompts used for a) Relation Extraction and b) filtering of uncertain sentences

**Figure 2:**
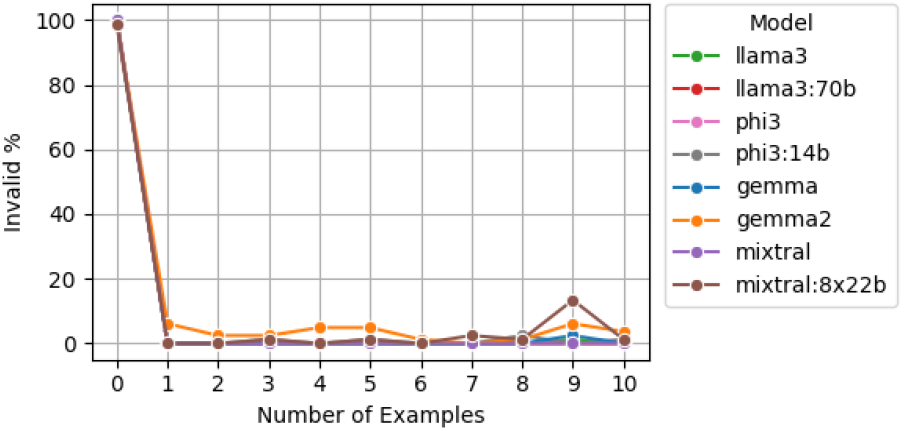
Analysis of the impact of the number of examples in the percentage of invalid LLM responses

The examples used in these analyses were generated by firstly querying Llama3 with two (one Positive and one Negative) manually crafted examples, using the final prompt, to obtain a large set of model generated responses. Then, the responses with correct predictions were manually curated into a set of ten final examples (five Positive and five Negative).

## 3. Results and Discussion

### 3.1. ncRNA-phenotype Relation Dataset

The obtained ncRNA-phenotype relation dataset successfully aggregates the five selected databases in an uniform format. On Figure 3, it is visible that there is small overlap between the databases, meaning that the inclusion of every database is pertinent. The attribution of unique identifiers to both ncRNAs and phenotypes guarantees the quality and non-ambiguity of the relations by forcing the rejection of wrongly registered, redundant or uninformative entries from the original databases. In Table 1, it is noticeable that the relations resulting from propagation to phenotype ancestors represent a great majority, meaning that this step significantly contributed to the completion of the dataset. However, in the ERL of phenotypes in disease descriptions, despite the high precision (0.973) of the method, some HPO Terms were wrongly attributed (statistically, about 3%), with this error being further amplified in the ancestor propagation. To mitigate this, a more restrictive smaller distance threshold could have been used, but in trade-off with a smaller number of disease descriptions with linked HPO Terms, and thus, relations.

**Table 1:**
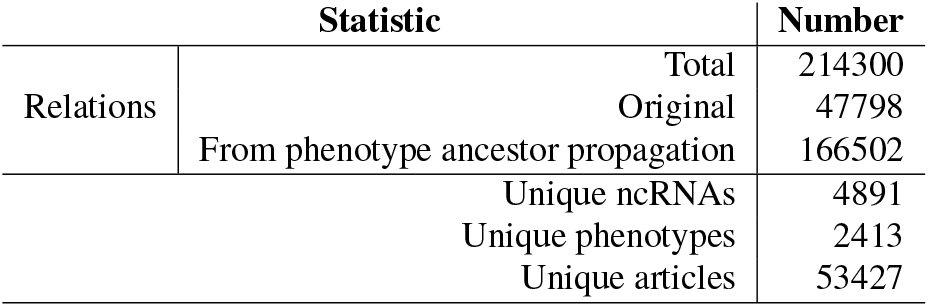
Statistics of the Relation Dataset

**Table 2:** Statistics for ncoRP

| Statistic |  | Number |
| --- | --- | --- |
| Annotated sentences |  | 208539 |
| Annotations | Total | 400557 |
|  | Positives | 84409 |
|  | Negatives | 316148 |
| Unique Relations | Total | 35295 |
|  | Positives | 3615 |
|  | Negatives | 34147 |
| Unique ncRNAs |  | 2835 |
| Unique phenotypes |  | 1118 |
| Annotated articles |  | 21608 |

**Table 3:** Evaluation results and metrics for ncoRP

| Evaluations |  |  |  |  | Metrics |  |  |  |  |
| --- | --- | --- | --- | --- | --- | --- | --- | --- | --- |
| TP | FP | FN | TN | Uncertain | Precision | Recall | F1-score | Uncertain % | Fleiss' Kappa |
| 54 | 17 | 57 | 14 | 38 | 0.761 | 0.468 | 0.593 | 21.1 | 0.431 |

**Figure 3:**
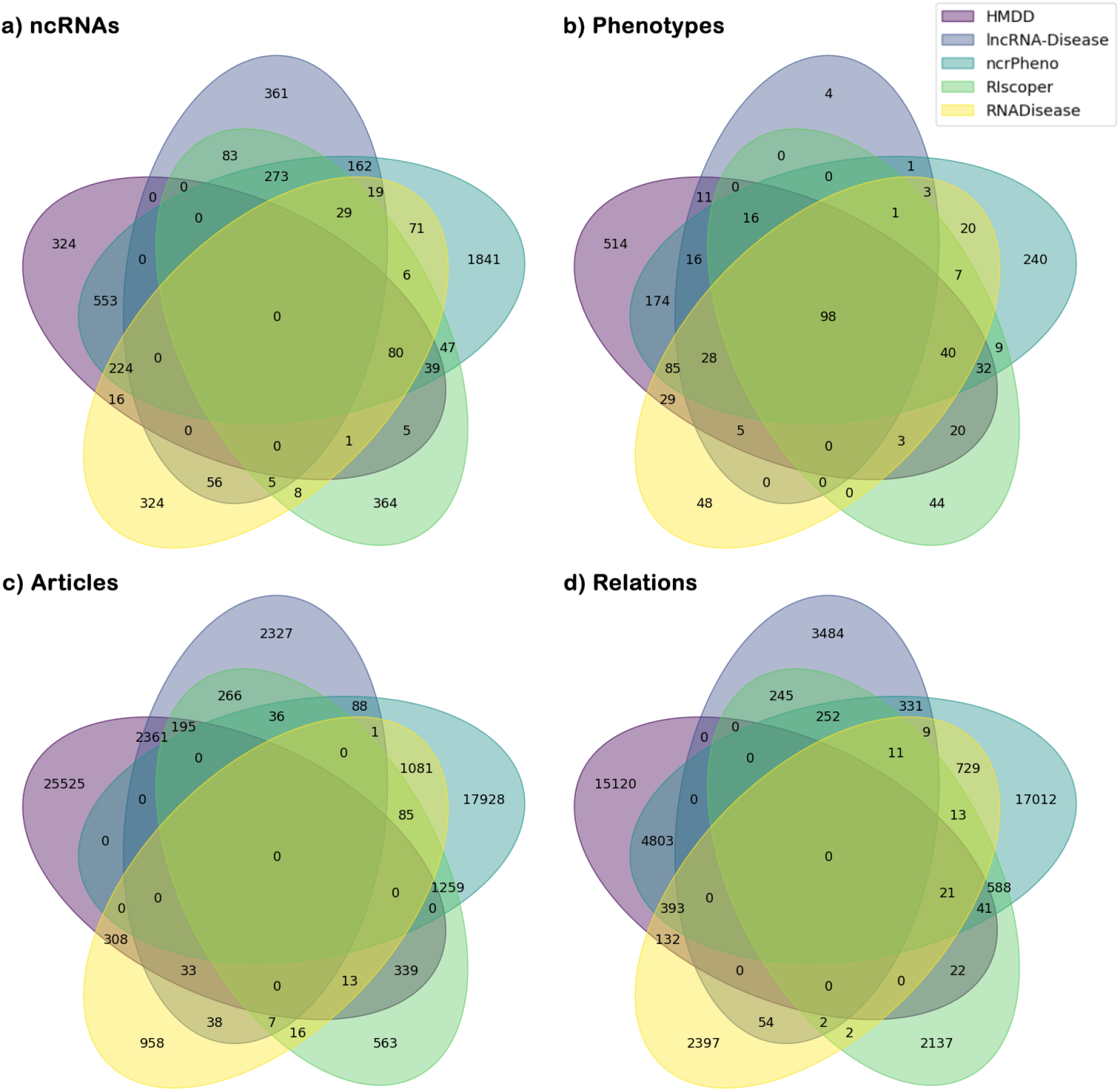
Overlap of the different databases in terms of a) unique ncRNAs, b) unique phenotypes, c) unique research articles and d) unique ncRNA-phenotype relations in the Relation Dataset

**Figure 4:**
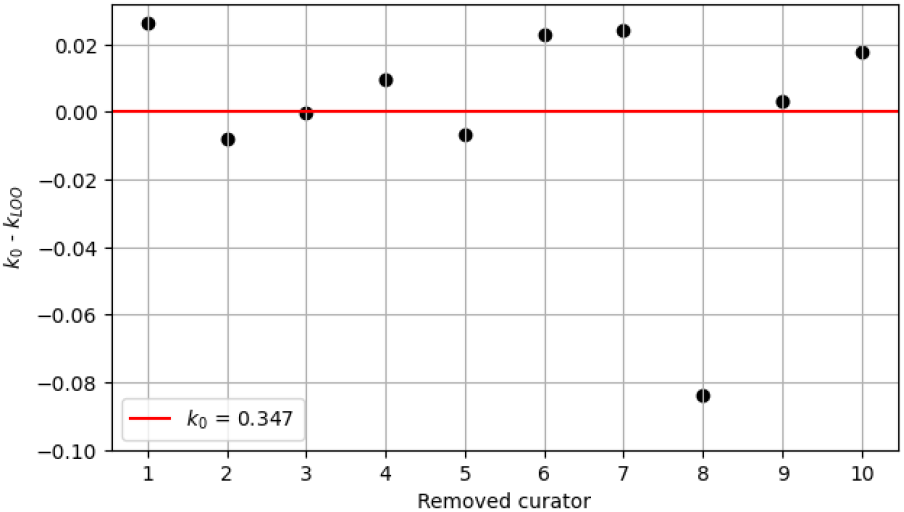
Differences between the baseline Fleiss’ Kappa and each Leave-One-Out Fleiss’ Kappa

**Figure 5:**
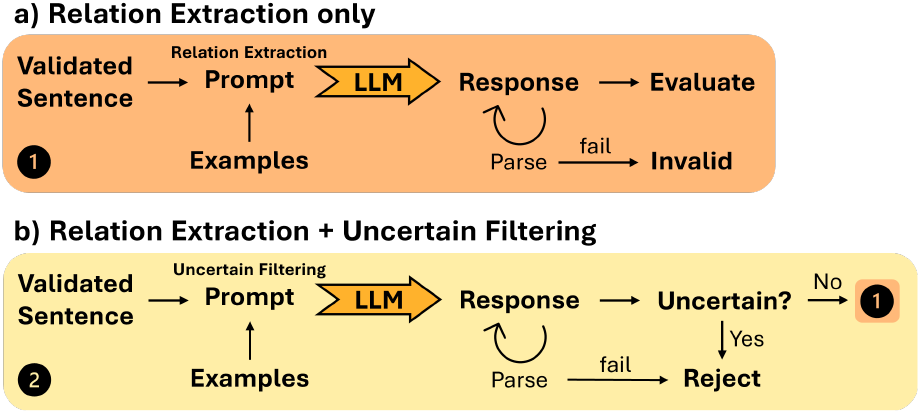
Visual representation of the Large Language Model Relation Extraction methodologies: a) Relation Extraction only and b) Relation Extraction preceded by uncertain sentence filtering

### 3.2. ncoRP

The produced ncRNA-phenotype corpus - ncoRP - contains more than 200,000 annotated sentences from a total of 21,608 scientific research articles concerning ncRNAs, with acceptable metrics (precision of 0.761 and F1-score of 0.593) considering the used method. However, based on the moderate inter-curator agreement (Fleiss’ Kappa of 0.431) and the fact that only about 0.1% of the corpus was evaluated, these metrics may not represent it fully, but are still a good representation. In the corpus, the great majority of sentences were labelled as *Negative*, as was expected with DSRE, which was also aggravated by the imposed article-specific restriction. Thus, by lifting this restriction, more *Positive* sentences could have been obtained, but possibly resulting in more FP instances. The obtained high FP and FN counts are mostly caused by the aforementioned problems inherent to DSRE and the high number of *Uncertain* sentences (21.1%) are caused by i) the method used to individualize sentences (resulting in some getting too long and complex to understand), ii) errors in the ERL of entities in the sentences (despite the filters used, some words were wrongly identified as entities that share the same name) and iii) that some sentences are in fact inherently uncertain in their meaning. Furthermore, this high number of *Uncertain* sentences, may have also hindered the observed inter-curator agreement.

### 3.3. LLM Relation Extraction

A sound method to obtain reliable uniform LLM RE responses was achieved by combining i) a prompt that instructed the model to produce a specific format (JSON), while including in-context examples with ii) a response parsing function, solving one of the main challenges of applying LLMs to close-ended tasks like RE. This enabled the automatic evaluation of responses with a ground-truth dataset (constructed from the validation results of ncoRP), which would have not been possible due to the diversity of possible responses these models normally generate. However, the employed process of prompt design could be improved as it consisted mainly of trial-and-error of different prompts, which made it difficult to comparatively evaluate their quality. As such, a more methodical approach should have been used.

The impact of in-context learning in LLM RE was studied by analysing the impact of the number of shots in performance (see Figure 6), with two main results arising. First, larger model sizes do not seem to increase performance, at least for the numbers of shots tested.

**Figure 6:**
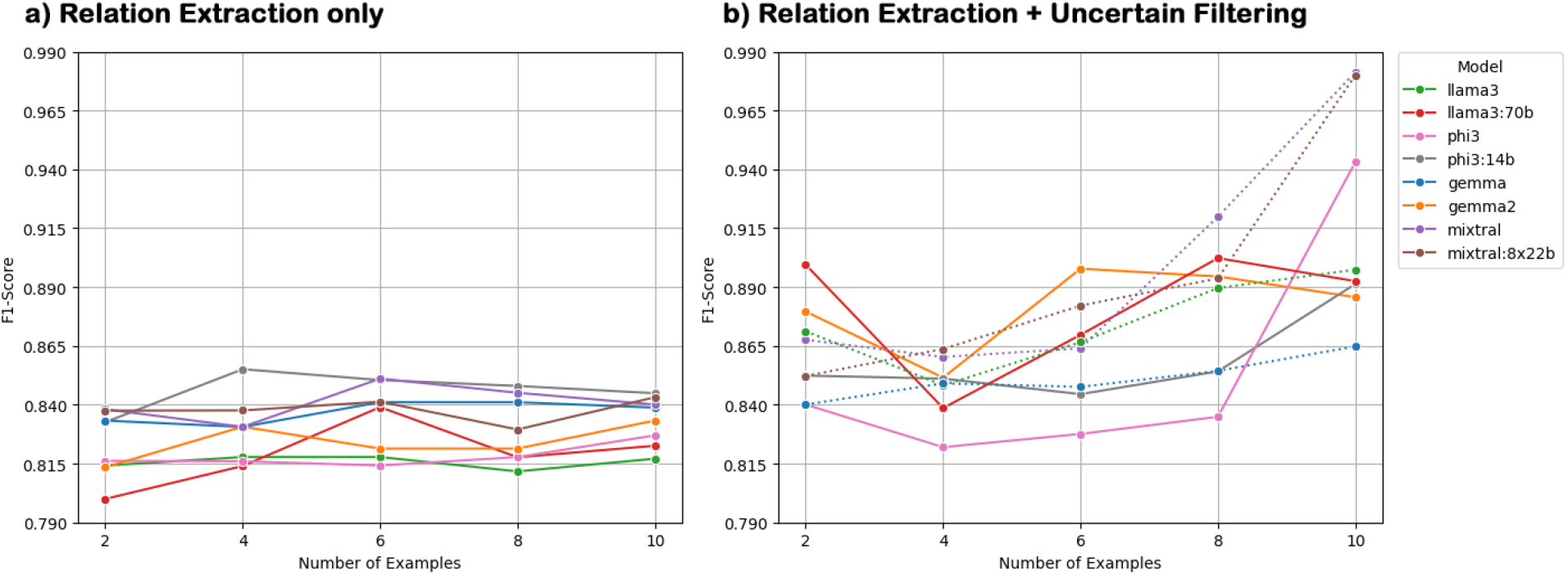
Large Language Model Relation Extraction performance analysis for different models and numbers of shots in a) a single Relation Extraction prompt and b) a Relation Extraction prompt preceded by an uncertain filtering prompt. Dotted lines represent models that resulted in more than 10% of invalid responses to the uncertain filtering prompt.

Perhaps better performance on large models could be achieved with larger numbers of shots, that would make use of their larger context windows. Second, the number of shots does not seem to have a clear impact on performance in the RE single prompt method, but higher numbers of shots seem to lead to better performance when combining the RE and uncertain filtering prompts. A possible explanation could be that more examples improve the performance of the uncertain filtering step, leading to more uncertain sentences being filtered, thus improving the performance in the RE step by making the model predict relations only in sentences it deems understandable. The results support this explanation in the sense that the best performing method also has the most percentage of sentences being filtered as uncertain. However, the obtained results could also be caused by the specific examples used and their order, and although it would be worthwhile to study different (possibly randomly selected/ordered) examples it would increase the complexity of the study beyond the scope of this work.

The best-found method combined the model Phi3 with ten shots, using the combined uncertain filtering and RE prompts, achieving an F1-score of 0.978, which outperforms RE state-of-the-art methods. However, a more thorough prompt design should have been applied for the uncertain sentence filtering prompt, as it resulted in a high number of invalid LLM responses in some models. Furthermore, the uncertain filtering prompt was only analysed in association with RE and its standalone performance (i.e. how good in fact it is at identifying uncertain sentences) was not fully examined as even manual analysis would be challenging due to the subjective nature of what constitutes an uncertain sentence. To conclude, despite the simplicity of the methods presented in this chapter, based mostly on prompting principles (namely in-context learning and CoT self-explanation) applied on small LLMs, very promising results were obtained, leaving very much apparent the potential of LLMs for RE, especially considering that these models do not require pre-existing labelled training data.

## 4. Conclusion

NcRNAs represent the majority of human gene products and are involved in a large set of important biological processes. Their dysregulation is associated with the origin and aggravation of various diseases, making them very relevant disease biomarkers and therapeutic agents. Despite their apparent importance, there is a lack of ncRNA functional annotation databases and information on these biomolecules is still sparsely distributed mainly in the form of scientific research articles. This coupled with the growing throughput of scientific text publications, makes it impossible to manually read and be aware of all the existing information.

This work demonstrates how AI can be employed, particularly through NLP techniques ranging different levels of complexity, to automatically extract and summarize the information available in these large textual datasets. First, a ncRNA-phenotype relation dataset was created, aggregating information from five ncRNA-disease functional annotation databases in a set of 214,300 unique relations. This enabled the creation of a ncRNA-phenotype relational corpus (ncoRP) through DSRE comprising 35,295 unique relations with a precision of 0.761 and a F1-score of 0.593. Both these contributions aim to mitigate the problem with the sparseness of information about ncRNAs. The developed pipelines can be easily adapted to be applied to general information extraction and summarization.

Currently, the AI scene is dominated by the LLM paradigm, for their complex understanding of written instructions and capacity to answer in coherent and meaningful generated text. In this work, the potential of LLMs for RE (and NLP tasks in general) is shown, giving insight on how these powerful models can be used beyond day-to-day tasks. As such, a LLM-based RE methodology was developed, making use of simple prompting principles, such as in-context learning and CoT self-explanation, to leverage LLM emergent capabilities. This method chains i) an uncertain sentence filtering prompt with a ii) RE prompt to predict if two entities share a relation in a sentence, yielding a very competent F1-score of 0.978, outperforming stateof-the-art deep learning RE methods, without requiring fine-tuning with large (often manually) labeled datasets. Despite the possibility of these results appearing overly optimistic (as they can be inflated due to the in-context examples used), they still express how powerful LLMs can be in helping to solve the aforementioned problems with ncRNA and general scientific information.

## 5. Future Work

This work shows the usefulness of text mining in information extraction and summarization, through its application on ncRNAs. As such, it would be insightful to apply the established pipelines to different biomedical entity pairs, such as “gene-drug” or “protein-protein”, or even going a step further with n-ary relations such as “gene-drug-tissue”. Furthermore, the development of a software tool capable of automatically annotating a given set of scientific research articles with relations involving any pair of entities stands as an interesting project.

The results obtained for LLMs, even employing simple prompting principles on small models, are already promising, as such, the application of more complex LLM approaches could further increase performance. Namely the use of i) larger models, ii) task-specific weight fine-tuning (which could be done using training data generated by these simpler approaches) or iii) refined in-context example generation/selection/ordering, are all topics that deserve further exploration.

## 6. Supplementary Information

All the code supporting this work is publicly available at https://github.com/davidcoscor/ncRNA-AI, including download URLs for the ncRNA-phenotype Relation Dataset and ncoRP.

## 7. Acknowledgements

We acknowledge FCT for funding through project “DeePer - Deep graph learning approaches to personalized medicine” (EXPL/CCI-BIO/0126/2021). This research was supported by Fundação para a Ciência e a Tecnologia through the fundings: UIDB/04046/2020 (DOI:10.54499/UIDB/04046/2020) and UIDP/04046/2020 (DOI:10.54499/UIDP/04046/2020)

Center grants from FCT, Portugal (to BioISI). Supported by FCT through funding of the LASIGE Research Unit, ref. UIDB/00408/2020 (https://doi.org/10.54499/UIDB/00408/2020) and ref. UIDP/00408/2020 (https://doi.org/10.54499/UIDP/00408/2020).

## References

[1] J. Niderla-Bielińska, E. Jankowska-Steifer, P. Włodarski, Non-coding rnas and human diseases: Current status and future perspectives, International Journal of Molecular Sciences 24 (2023). doi:10.3390/ijms241411679.

[2] J. Zhang, S. Zou, L. Deng, Gene ontology-based function prediction of long non-coding rnas using bi-random walk 06 biological sciences 0601 biochemistry and cell biology, BMC Medical Genomics 11 (2018). doi:10.1186/s12920-018-0414-2.

[3] L. Sang, L. Yang, Q. Ge, S. Xie, T. Zhou, A. Lin, Subcellular distribution, localization, and function of noncoding rnas, Wiley Interdisciplinary Reviews: RNA 13 (2022) e1729.

[4] T. Loganathan, G. P. D. C, Non-coding rnas in human health and disease: potential function as biomarkers and therapeutic targets, Functional and Integrative Genomics 23 (2023). doi:10.1007/s10142-022-00947-4.

[5] E. López-Jiménez, E. Andrés-León, The implications of ncrnas in the development of human diseases, Non-coding RNA 7 (2021) 17.

[6] D. F. Sousa, F. M. Couto, K-ret: knowledgeable biomedical relation extraction system, Bioinformatics 39 (2023). doi:10.1093/bioinformatics/btad174.

[7] A. Lamurias, L. A. Clarke, F. M. Couto, Extracting microrna-gene relations from biomedical literature using distant supervision, PLoS ONE 12 (2017). doi:10.1371/journal.pone.0171929.

[8] D. Khurana, A. Koli, K. Khatter, S. Singh, Natural language processing: state of the art, current trends and challenges, Multimedia Tools and Applications 82 (2023). doi:10.1007/s11042-022-13428-4.

[9] K. Detroja, C. K. Bhensdadia, B. S. Bhatt, A survey on relation extraction, Intelligent Systems with Applications 19 (2023). doi:10.1016/j.iswa.2023.200244.

[10] D. Sousa, A. Lamurias, F. M. Couto, A silver standard corpus of human phenotype-gene relations, Proceedings of the 2019 Conference of the North American Chapter of the Association for Computational Linguistics: Human Language Technologies 1 (2019). URL: http://arxiv.org/abs/1903.10728 http://dx.doi.org/10.18653/v1/N19-1152. doi:10.18653/v1/N19-1152.

[11] W. X. Zhao, K. Zhou, J. Li, T. Tang, X. Wang, Y. Hou, Y. Min, B. Zhang, J. Zhang, Z. Dong, Y. Du, C. Yang, Y. Chen, Z. Chen, J. Jiang, R. Ren, Y. Li, X. Tang, Z. Liu, P. Liu, J.-Y. Nie, J.-R. Wen, A survey of large language models, 2023. URL: http://arxiv.org/abs/2303.18223.

[12] C. Cui, B. Zhong, R. Fan, Q. Cui, Hmdd v4.0:ã database for experimentally supported human microrna-diseaseãssociations, Nucleic Acids Research 52 (2024) D1327–D1332. doi:10.1093/nar/gkad717.

[13] X. Lin, Y. Lu, C. Zhang, Q. Cui, Y. D. Tang, X. Ji, C. Cui, Lncrnadisease v3.0: an updated database of long non-coding rna-associated diseases, Nucleic Acids Research 52 (2024) D1365– D1369. doi:10.1093/nar/gkad828.

[14] W. Zhang, G. Yao, J. Wang, M. Yang, J. Wang, H. Zhang, W. Li, ncrpheno: a comprehensive database platform for identification and validation of disease related noncoding rnas, RNA Biology 17 (2020) 943–955. doi:10.1080/15476286.2020.1737441.

[15] H. Zheng, L. Xu, H. Xie, J. Xie, Y. Ma, Y. Hu, L. Wu, J. Chen, M. Wang, Y. Yi, Y. Huang, D. Wang, Riscoper 2.0: A deep learning tool to extract rna biomedical relation sentences from literature, Computational and Structural Biotechnology Journal 23 (2024) 1469–1476. doi:10.1016/j.csbj.2024.03.017.

[16] J. Chen, J. Lin, Y. Hu, M. Ye, L. Yao, L. Wu, W. Zhang, M. Wang, T. Deng, F. Guo, Y. Huang, B. Zhu, D. Wang, Rnadisease v4.0: an updated resource of rna-associated diseases, providing rna-disease analysis, enrichment and prediction, Nucleic Acids Research 51 (2023) D1397–D1404. doi:10.1093/nar/gkac814.

[17] N. Reimers, I. Gurevych, Sentence-bert: Sentence embeddings using siamese bert-networks, in: Proceedings of the 2019 Conference on Empirical Methods in Natural Language Processing, Association for Computational Linguistics, 2019. URL: https://arxiv.org/abs/1908.10084.

[18] J. Johnson, M. Douze, H. Jégou, Billion-scale similarity search with GPUs, IEEE Transactions on Big Data 7 (2019) 535–547.

[19] M. A. Gargano, N. Matentzoglu, B. Coleman, E. B. Addo-Lartey, A. V. Anagnostopoulos, J. Anderton, P. Avillach, A. M. Bagley, E. Bakštein, J. P. Balhoff, G. Baynam, S. M. Bello, M. Berk, H. Bertram, S. Bishop, H. Blau, D. F. Bodenstein, P. Botas, K. Boztug, J. Čady, T. J. Callahan, R. Cameron, S. J. Carbon, F. Castellanos, J. H. Caufield, L. E. Chan, C. G. Chute, J. Cruz-Rojo, N. Dahan-Oliel, J. R. Davids, M. Dieuleveult, V. Souza, B. B. de Vries, E. Vries, J. R. DePaulo, B. Derfalvi, F. Dhombres, C. Diaz-Byrd, A. J. Dingemans, B. Donadille, M. Duyzend, R. Elfeky, S. Essaid, C. Fabrizzi, G. Fico, H. V. Firth, Y. Freudenberg-Hua, J. M. Fullerton, D. L. Gabriel, K. Gilmour, J. Giordano, F. S. Goes, R. G. Moses, I. Green, M. Griese, T. Groza, W. Gu, J. Guthrie, B. Gyori, A. Hamosh, M. Hanauer, K. Hanušová, Y. He, H. Hegde, I. Helbig, K. Holasová, C. T. Hoyt, S. Huang, E. Hurwitz, J. O. Jacobsen, X. Jiang, L. Joseph, K. Keramatian, B. King, K. Knoflach, D. A. Koolen, M. L. Kraus, C. Kroll, M. Kusters, M. S. Ladewig, D. Lagorce, M. C. Lai, P. Lapunzina, B. Laraway, D. Lewis-Smith, X. Li, C. Lucano, M. Majd, M. L. Marazita, V. Martinez-Glez, T. H. McHenry, M. G. McInnis, J. A. McMurry, M. Mihulová, C. E. Millett, P. B. Mitchell, V. Moslerová, K. Narutomi, S. Nematollahi, J. Nevado, A. A. Nierenberg, N.N. Čajbiková, J. I. Nurnberger, S. Ogishima, D. Olson, A. Ortiz, H. Pachajoa, G. P. Nanclares, A. Peters, T. Putman, C. K. Rapp, A. Rath, J. Reese, L. Rekerle, A. M. Roberts, S. Roy, S. J. Sanders, C. Schuetz, E. C. Schulte, T. G. Schulze, M. Schwarz, K. Scott, D. Seelow, B. Seitz, Y. Shen, M. N. Similuk, E. S. Simon, B. Singh, D. Smedley, C. L. Smith, J. T. Smolinsky, S. Sperry, E. Stafford, R. Stefancsik, R. Steinhaus, R. Strawbridge, J. C. Sundaramurthi, P. Talapova, J. A. Castano, P. Tesner, R. H. Thomas, A. Thurm, M. Turnovec, M. E. van Gijn, N. A. Vasilevsky, M. VlČková, A. Walden, K. Wang, R. Wapner, J. S. Ware, A. A. Wiafe, S. A. Wiafe, L. D. Wiggins, A. E. Williams, C. Wu, M. J. Wyrwoll, H. Xiong, N. Yalin, Y. Yamamoto, L. N. Yatham, A. K. Yocum, A. H. Young, Z. Yüksel, P. P. Zandi, A. Zankl, I. Zarante, M. Zvolský, S. Toro, L. C. Carmody, N. L. Harris, M. C. Munoz-Torres, D. Danis, C. J. Mungall, S. Köhler, M. A. Haendel, P. N. Robinson, The human phenotype ontology in 2024: phenotypes around the world, Nucleic Acids Research 52 (2024) D1333–D1346. doi:10.1093/nar/gkad1005.

[20] B. A. Sweeney, A. I. Petrov, C. E. Ribas, R. D. Finn, A. Bateman, M. Szymanski, W. M. Karlowski, S. E. Seemann, J. Gorodkin, J. J. Cannone, R. R. Gutell, S. Kay, S. Marygold, G. D. Santos, A. Frankish, J. M. Mudge, R. Barshir, S. Fishilevich, P. P. Chan, T. M. Lowe, R. Seal, E. Bruford, S. Panni, P. Porras, D. Karagkouni, A. G. Hatzigeorgiou, L. Ma, Z. Zhang, P. J. Volders, P. Mestdagh, S. Griffiths-Jones, B. Fromm, K. J. Peterson, I. Kalvari, E. P. Nawrocki, A. S. Petrov, S. Weng, P. Bouchard-Bourelle, M. Scott, L. M. Lui, D. Hoksza, R. C. Lovering, B. Kramarz, P. Mani, S. Ramachandran, Z. Weinberg, Rnacentral 2021: Secondary structure integration, improved sequence search and new member databases, Nucleic Acids Research 49 (2021) D212–D220. doi:10.1093/nar/gkaa921.

[21] F. M. Couto, A. Lamurias, Mer: A shell script and annotation server for minimal named entity recognition and linking, Journal of Cheminformatics 10 (2018). doi:10.1186/s13321-018-0312-9.

[22] Şenay Kafkas, S. Althubaiti, G. V. Gkoutos, R. Hoehndorf, P. N. Schofield, Linking common human diseases to their phenotypes; development of a resource for human phenomics, Journal of Biomedical Semantics 12 (2021). doi:10.1186/s13326-021-00249-x.

[23] J. Zhao, Z. Yao, Z. Yang, H. Yu, Self-explain: Teaching large language models to reason complex questions by themselves, 2023.

